# msVolcano: a flexible web application for visualizing quantitative proteomics data

**DOI:** 10.1101/038356

**Authors:** Sukhdeep Singh, Marco Y. Hein, A. Francis Stewart

## Abstract

We introduce msVolcano, a web application, for the visualization of label-free mass spectrometric data. It is optimized for the output of the MaxQuant data analysis pipeline of interactomics experiments and generates volcano plots with lists of interacting proteins. The user can optimize the cutoff values to find meaningful significant interactors for the tagged protein of interest. Optionally, stoichiometries of interacting proteins can be calculated. Several customization options are provided to the user for flexibility and publication-quality outputs can also be downloaded (tabular and graphical).

Availability: msVolcano is implemented in R Statistical language using Shiny and is hosted at server in-house. It can be accessed freely from anywhere at http://projects.biotec.tudresden.de/msVolcano/

## 1 Introduction

The analysis of protein-protein interactions and complex networks using affinity purification or affinity enrichment and mass spectrometry (AP/MS,AE/MS) is one of the most commonly used applications in proteomics. The technology produces high quality protein interaction data [1] and is scalable to proteome-wide levels [2]. Even though isotope labeling methods have been developed to detect and quantify protein-protein interactions [3], label-free approaches are gaining momentum due to their simplicity and applicability [4]. While different quantification strategies exist for label-free data, such as those based on spectral counting, methods that make use of peptide intensities (also known as extracted ion currents) are regarded as the most accurate [5, 6]. Such methods generate quantitative profiles of peptides or proteins across samples, which can be analyzed by established statistical methods, e.g. by a modified t-test across replicate experiments [7].

MaxQuant is an integrated suite of algorithms for the analysis of high-resolution quantitative MS data [8]. Its MaxLFQ module normalizes the contribution of individual peptide fractions and extracts the maximum available quantitative information to calculate highly reliable relative label-free quantification (LFQ) intensity profiles [6], which are exported as tab-limited text files for the downstream analysis.

To identify interactors of a tagged protein of interest (termed the ‘bait’), in the presence of a vast number of background binding proteins, replicates of affinity-enriched bait samples are compared to a set of negative control samples. A student’s t-test or Welch’s test can be used to determine those proteins that are significantly enriched along with the specific baits. A volcano plot is a good way to visualize this kind of analysis [9]. When the resulting differences between the logarithmized mean protein intensities between bait and the control samples are plotted against the negative logarithmic p values derived from the statistical test, unspecific background binders center around zero. The enriched interactors appear on the right section of the plot, whereas ideally no proteins should appear on the left section when compared to an empty control (because these would represent proteins depleted by the bait). The higher the difference between the group means (i.e. the enrichment) and the p value (i.e. the reproducibility), the more the interactors shift towards the upper right section of the plot, which represents the area of highest confidence for a true interaction.

Though label-free methods are nowadays as accurate as the isotope-based methods, false positives may appear enriched alongside the true positive interaction partners [10]. Identifying background proteins and defining a threshold that separates these from true interactors is a critical step during data analysis, and often benefits from some manual optimization.

## 2 Description

To facilitate the analysis and presentation of AE-MS data, we present msVolcano, which is a user modulated, freely accessible web application. It requires the MaxQuant output of an interaction dataset that was analysed using the MaxLFQ module. LFQ intensity profiles retain the absolute scale from the original summed- up peptide intensities [6], serving as a proxy for absolute protein abundance. The purpose of msVolcano is to implement all steps of downstream data analysis into a simple and intuitive user interface that requires no bioinformatics knowledge or specialized software. To this end, msVolcano automatically extracts relevant data columns, filters out hits to the decoy database and potential contaminants and imputes missing values by simulated noise. A visual Quality Control (QC) output is generated allowing the user to monitor the correlation between replicates, fraction of missing values and behaviour of the population of imputed values.

A user-defined statistical test is then performed between selected bait and control samples and the tool generates a volcano plot. We implemented a recently introduced hyperbolic curve threshold [11], based on the given formula

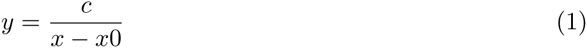

where *c* = curvature, x0 = minimum fold change, thus dividing enriched proteins into mildly and strongly enriched [11]. The cutoff parameters can be adjusted by the user and monitored by the graphical output. The user has access to the plot aesthetics and can view the original input file and its subset for significant interactors in the inbuilt browser. A publication-quality PDF plot can be generated and exported along with the subset of original data limited to the significant interactors. Next to the identities of interacting proteins, their stoichiometries relative to their bait is crucial for the understanding of the molecular function of protein complexes [2, 12]. Thus, optional stoichiometry calculations have been implemented in the code. We use a modified version of intensity-based absolute quantification (iBAQ) [13] for the estimation of protein abundance for stoichiometry calculations, where LFQ intensities are normalized by the number of theoretical tryptic peptides between 7 and 30 amino acids, as described [2](Fig 1b). Theoretical peptides are pre-calculated for the most commonly used proteomes of model organisms and are matched based on the proteins’ uniprot IDs. Stoichiometry calculations are based on the given formula

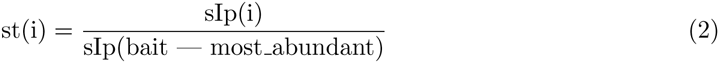

where

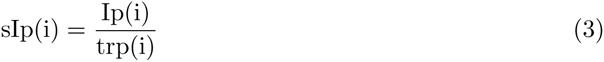

where

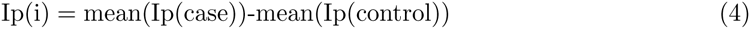

where st = stoichiometry, sip = size normalised protein intensity, Ip = protein intensity, trp = number of theoretical peptides of protein

**Figure 1:**
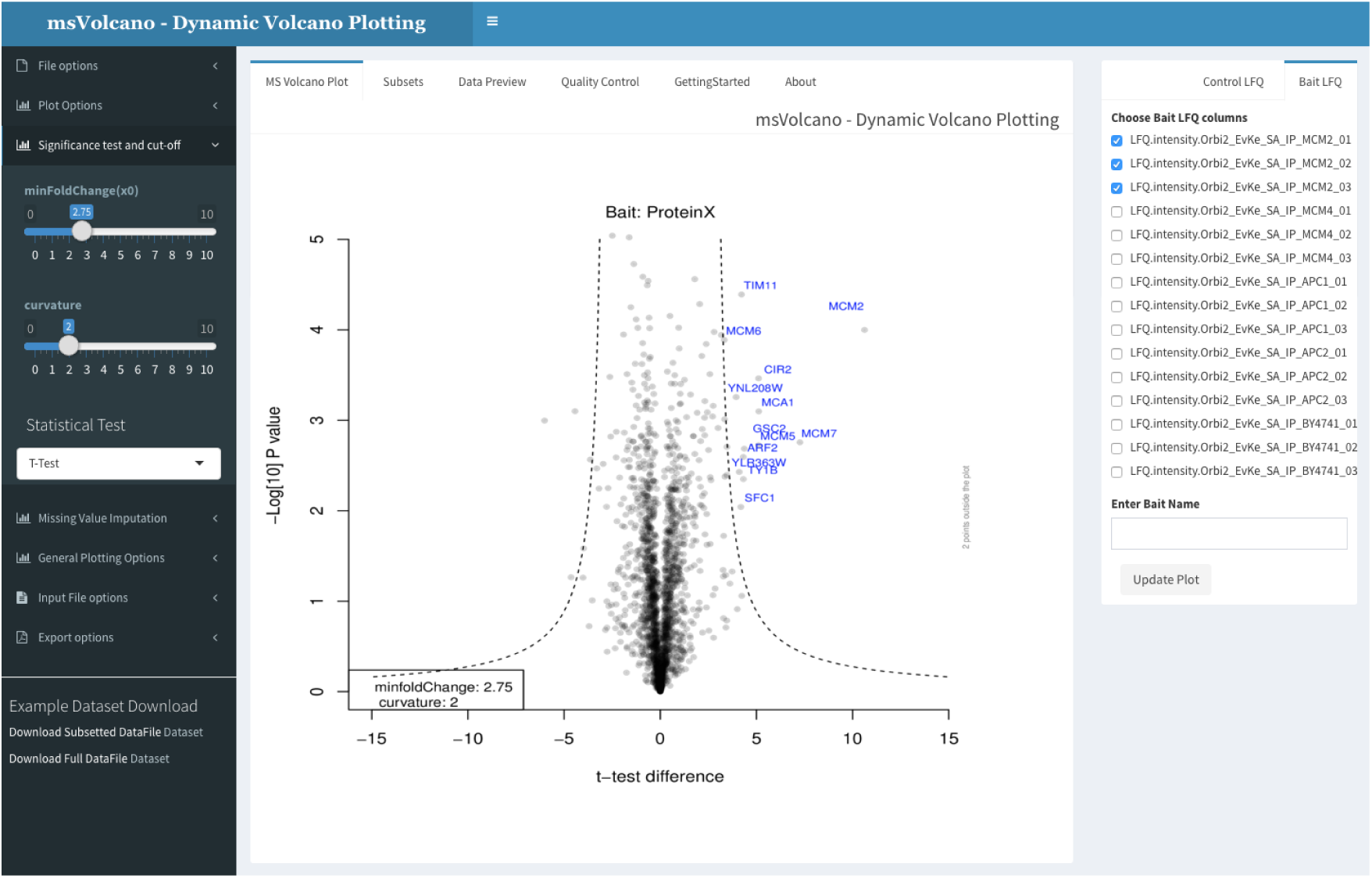
The interface is divided into three sections, sidebar, body panel and column selection panel (left to right). Sidebar provides an access to the file upload, plot aesthetics, cutoff parameters, missing data imputation, stoichiometry and the export options. The body panel has five different tabs, where the default panel labeled as “MS Volcano Plot” displays the volcano plot. Second tab, “Subsets” displays the filtered input data for the significant interactors. “Data Preview” tab displays the user inputted data for scrutiny, “GettingStarted” and “About” tab display the specific and general information about the web interface. When user uploads a file or enters a ftp link, all LFQ columns are scanned and displayed in the column selection tab on the right side. User now selects respective bait and control columns (minimum two) and optionally enters the name of bait in the provided text box. As the ‘Update Plot’ button is pressed, the plot is generated simultaneously.

**Figure 2:**
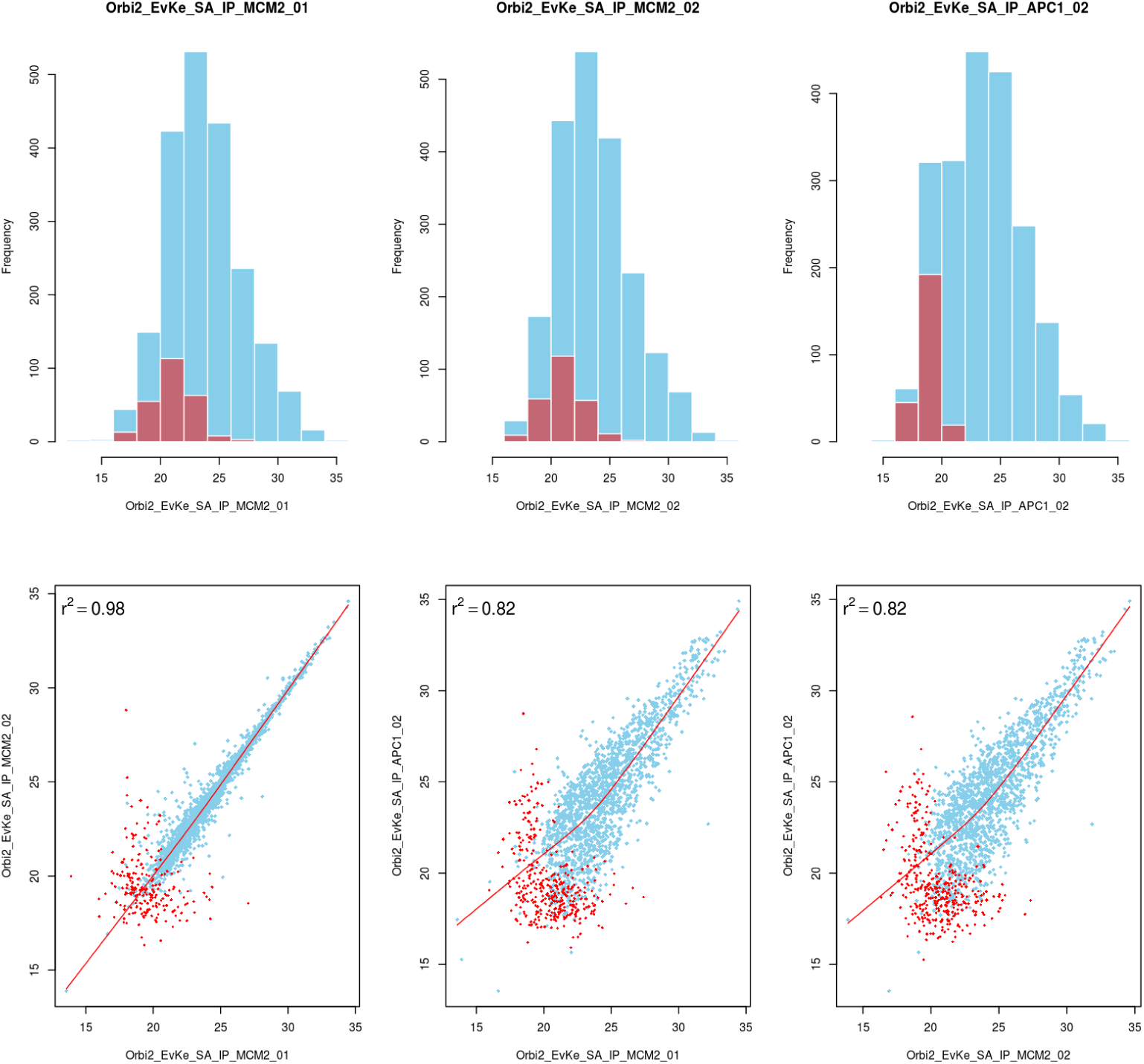
Default QC plot using a dataset from budding yeast study (sample data in msVolcano) [11] A) top row displaying the distribution of the raw values (LFQ intensites - in blue) overlayed with the distribution of imputed values (in red) per LFQ column selected B) 2x2 scatter plots between the chosen LFQ columns with local regression(lowess) displayed as a red line with pearson’s correlations coefficient. For the visual aestheticity, the number of scatter plots are restricted to the number of histograms displayed above them.

## 3 Conclusion

msVolcano provides a web-platform for the quick visualization of label-free mass spectrometric data and can be freely accessed globally. With the underlying hyberbolic curve parameters and other statistics, user can intuitively isolate the true protein interaction partners from the false positives, without the need of writing a code. With its ftp file input support, the user can quickly analyse and re-analyse the results of the interactomics experiment present on their own cloud servers and along with the calculated optional stoichiometries, all the results can be exported in publication quality tabular or graphical format.

## Funding

This work was supported by the EU 7th Framework integrated project, SyBoSS (www.syboss.eu).

